# Segmentation and analysis of mother machine data: SAM

**DOI:** 10.1101/2020.10.01.322685

**Authors:** Deb Sankar Banerjee, Godwin Stephenson, Suman G. Das

**Affiliations:** Department of Physics, Carnegie Mellon University, Pittsburgh, PA 15213, USA; Simons Centre for the Study of Living Machines, National Centre for Biological Sciences, Bangalore 560065, India; Institute for Biological Physics, University of Cologne, Cologne, Germany

## Abstract

Time-lapse imaging of bacteria growing in micro-channels in a controlled environment has been instrumental in studying the single cell dynamics of bacterial growth. This kind of a microfluidic setup with growth chambers is popularly known as *mother machine* [1]. In a typical experiment with such a set-up, bacterial growth can be studied for numerous generations with high resolution and temporal precision using image processing. However, as in any other experiment involving imaging, the image data from a typical mother machine experiment has considerable intensity fluctuations, cell intrusion, cell overlapping, filamentation etc. The large amount of data produced in such experiments makes it hard for manual analysis and correction of such unwanted aberrations. We have developed a modular code for segmentation and analysis of mother machine data (SAM) for rod shaped bacteria where we can detect such aberrations and correctly treat them without manual supervision. We track cumulative cell size and use an adaptive segmentation method to avoid faulty detection of cell division. SAM is currently written and compiled using MATLAB. It is fast (∼ 15 *min/GB* of image) and can be efficiently coupled with shell scripting to process large amount of data with systematic creation of output file structures and graphical results. It has been tested for many different experimental data and is publicly available in Github.

## I. INTRODUCTION

An integral and almost inseparable part of modern image analysis is segmentation of an image into meaningful objects or regions of interest. Though image segmentation has been studied for long it still presents with new challenges in analysis and often times a predetermined set of operations produces a poor quality of segmentation due to the noise and variability in biological images. Many studies have developed image analysis methods for bacterial growth analysis from mother machine data [2–11] using conventional and machine learning methods. Here we present a set of modular programs which can take a stack of images of rod shaped bacteria from a mother machine experiment and segment them to produce an easy to read data structure with the information of cell divisions. This data structure can be used to track all cells of four consecutive generations and hundreds of divisions of the mother cell (the cell at the end of the channel).

We first introduce the main working principle of SAM and give a brief overview of how it can be used to analyze and extract cellular information from mother machine image data. We present some sample results of cell division statistics using the cell data obtained from SAM. The code is publicly available in Github with sufficient documentation. We also provide additional methods developed to easily handle the structured cell data extracted by SAM. The main routines being written in a higher level language, are easy to follow and can be modified to suit the specific need of a study. We hope our simple image segmentation workflow can be understood and used by anyone with entry level knowledge of MATLAB. We have provided a separate, simpler and more documented Github training repository for beginners.

## II. AIM OF THE IMAGE ANALYSIS

We have developed the image processing workflow to detect cell divisions and track the cell lineages from fluorescence images obtained from a mother machine experiment (Fig. 1a). The channels we analyze are open at one end only (Fig. 1b) and the cells exit the channel from the open end as they divide and fill the channel. The cell at the closed end, referred as *old-pole mother cell* keeps dividing and can be tracked for many divisions. The other cells grow and divide as well but eventually exit the channel after few rounds of division of the old-pole cell. The major difficulties in the analysis of such data come from experimental aberrations such as large fluctuations in intensity, cell intrusion in the channel, cell overlapping and sticky cells at the channel end. We have tried to eliminate such aberrations without manual input in our workflow. We have employed tracking of various individual and collective features of the cells inside the channel to correctly capture cell division events and to avoid erratic detection of cell division due to intensity fluctuations and overlapping (which may lead to “rejoining” of newly divided cells if not corrected for) in newborn cells. In the cell data structure each cell of a channel has an unique identity number and we keep track of the parent cells which enables us to track cell divisions of all the cells up to four successive generations (where the total population consist of 8 cells and that almost fills the channel so further tracking of all 8 divisions becomes impossible due to extrusion of cells).

**FIG. 1.**
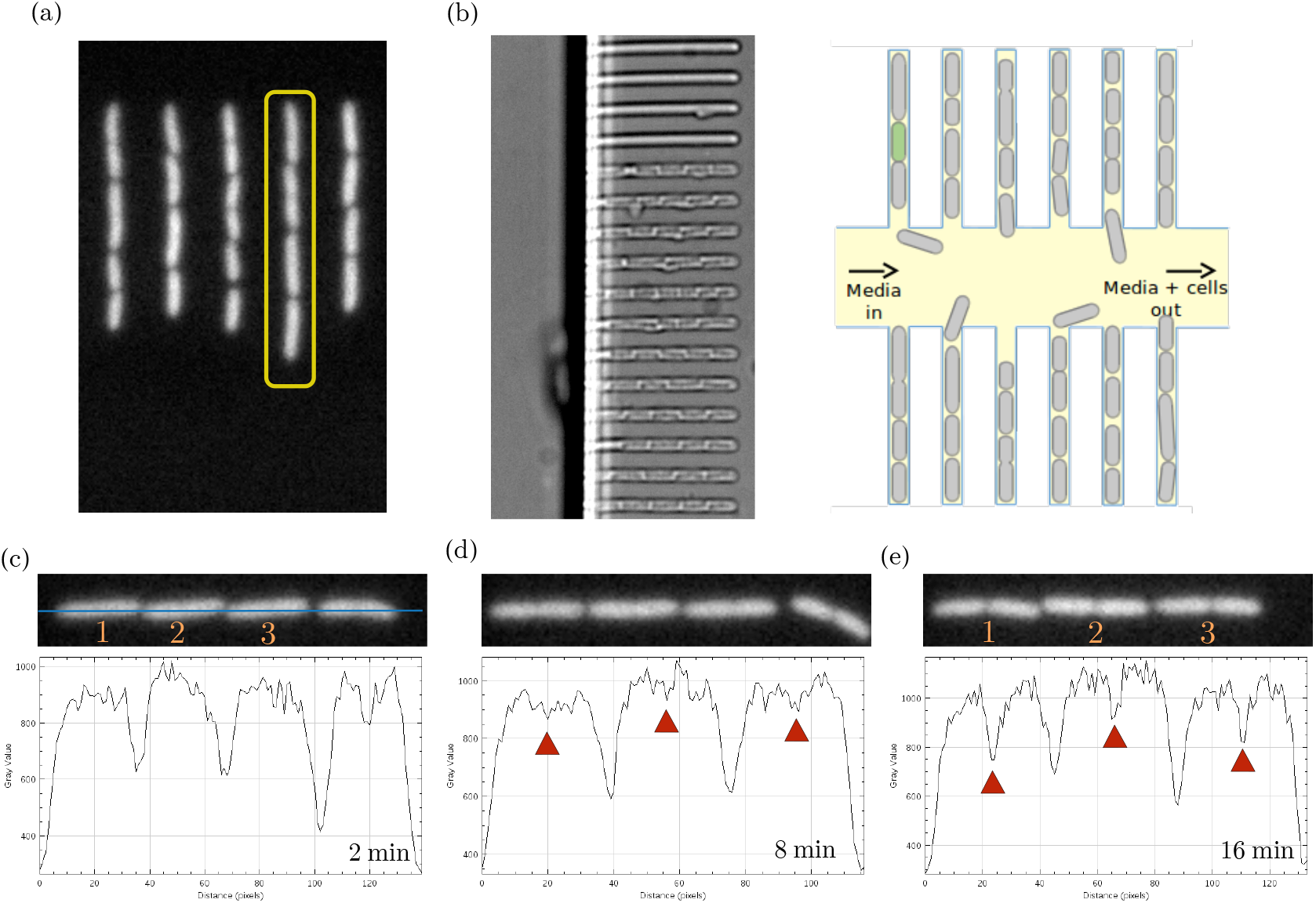
Features of mother machine data: (a) Typical mother machine image data shows florescently tagged *E. Coli* cells growing in parallel chambers/channels. We separate single channels (e.g., shown in yellow box) and save them as image stacks (.tif files) to be analyzed by SAM. (b) A phase contrast snapshot of the mother machine growth chambers (left) show bacteria cells inside the channels. A schematic diagram (right) shows the channels have the bottom end open for nutrient inflow. Bacteria cells keep growing and dividing inside the channel and as their numbers increase the cells in the bottom of the channel move out and get removed (cell extrusion) by the media flow in the device. (c-e) The midline (in blue) projection of intensity shows the large intensity dips in between two cells. Small intensity minima start developing when cells start to divide (marked in red). The main objective of image segemntation here is to correctly capture and categorize the new emerging minima to detect cell division events.

The main feature that helps in detecting cell division is the increasing intensity dip at the middle of a large cell. This dip signifies an approaching cell division (Fig. 1c-d). We define a threshold prominence (a relative strength) for the minimum to declare that a cell has divided. Cell to cell variability in intensity values makes it difficult to define global threshold values (Fig. 1e). A naive threshold based segmentation fails to detect cell divisions correctly because of this intensity variation.

### III. WORKING PRINCIPLE

Here we discuss the main algorithm and the major steps involved in the workflow employed in SAM. The entire process of data analysis starting from the raw unprocessed image data from mother machine to statistics of cell division can be described in three major steps:

1. Pre-processing the whole image of the microfluidic device: We break down the image into small image stacks each containing a single channel. This step is manually done in the current version of the workflow. We hope to include an automatic breakdown of the image in a later version.
2. Segmentation and analysis of channel data set: We use SAM for detection of individual cells and tracking of cell divisions from the whole set of channels. During tracking all common experimental noise and aberrations are automatically removed or corrected for without any manual intervention. All cell division and extrusion events are recorded in a lineage tractable data structure.
3. Further analysis and visualization: We have developed a set of codes to enable easy analysis of the structured cell data and visualization of the cell division statistics.

The second step (this is basically the step done by SAM) consists of many different operations on the image stack ending with producing a detailed data structure for cell division events for each channel. A short breakdown of the major operations performed in SAM (Fig. 2a) is given below:

**FIG. 2.**
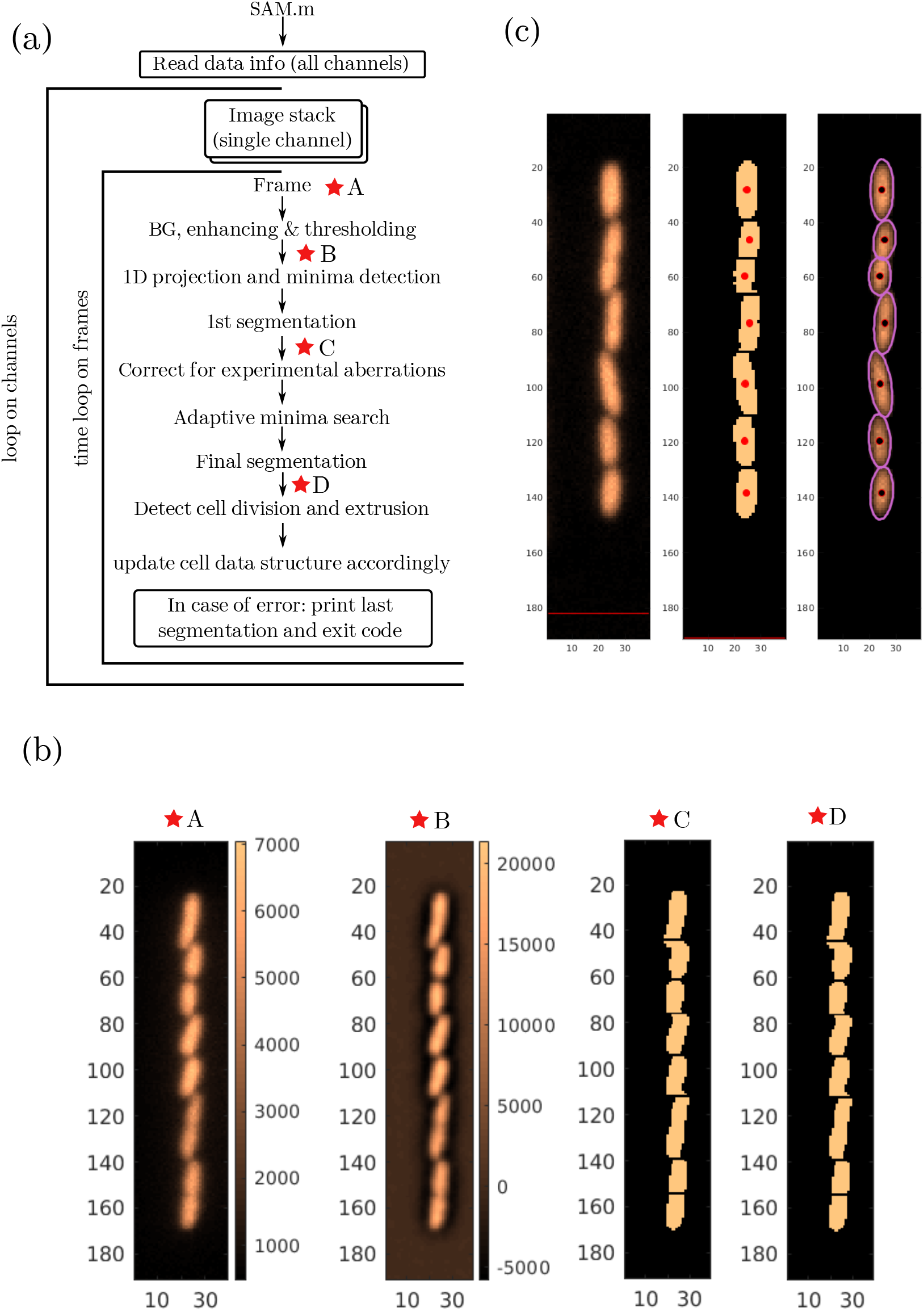
Segmentation and cell detection: (a) A reduced flowchart of the main working principle of SAM. We employ repeated segmentation based on a trial and error evaluation of segmentation quality. (b) The four frames show the images at the four star marked points within the flowchart. This particular frame being devoid of aberrations the two segmentations are very similar. (c) A sample SAM output figure shows segmentation of a frame where the raw image (left) was segmented and binarized (middle) to detect the individual cells and an ellipse was fitted to each cell (right). The black dot marks the centroid of the cell.

- All the channels (image stack) saved as .tif files in a particular directory (named “im” here) are read one by one in a data loop as image stack data. This image stack data will be used in the analysis of each frame.
- The frames saved in the 3D image stack data (named “FinalImage”) are called in the time loop. Each frame was processed and enhanced. Then a mid-line (long axis of the channel, see Fig. 1c) intensity projection was calculated by averaging over the width of the channel. The intensity minima were then detected from this one dimensional data.
- The first segmentation was performed (Fig. 2b) after this.
- The segmentation was done in two steps- a binary image of the frame with only the cells (pixel inside cell: High, pixel outside cell: Low) was created then this binary image was cut into separate connected objects (high value pixels that are adjacent) using the minima positions such that each connected objects correspond to one cell. These connected objects were saved in a structured array (named “CC”) for further analysis.
- This first segmentation is used in various consistency cross-checks to validate the segmentation. When aberrations or inconsistencies were detected different specialized correction or adaptive detection operations were run in a bid to get rid of the aberrations or inconsistencies.
- A final segmentation was performed after all corrections and the cells were saved in a structured array (named “CC2”) for further analysis.
- A measure of cell size was estimated from the major axis length of a fitted ellipse on each cell (Fig. 2c).
- In the case of a cell division and/or cell extrusion event the cell data (named “cell id”) was updated accordingly.
- These operations were repeated on each frame for each channel to extract cell division data from each channel.

The segmentation needs some user defined input parameters. All such parameters are to be defined in the main code “SAM.m”. The major quantities that have to be judiciously chosen are: (i) A global threshold (named “thr”) to create binary images. As a simple approximation an average background intensity can be set as this threshold. (ii) A rough estimate of typical cell size in pixels at birth and at division. Few parameters (such as minimum possible cell size, minimum distance between intensity minima etc) has to be determined from this estimate. These two quantities may depend on cell type, growth condition and microscopy settings.

## IV. DATA STRUCTURE AND CELL DIVISION STATISTICS

Here we discuss the output data structure and sample cell division statistics obtained using SAM. In all the presented analysis here we used SAM to analyze the publicly available [time lapse image data](https://figshare.com/collections/Data_from_long-term_growth_data_of_Escherichia_coli_at_a_single-cell_level/3493548) for *E. coli* cells from the study of Tanouchi *et al* [12, 13].

The output of SAM, a cell data created to record cell division events records six essential information for each cell. These six information are time of birth (Tbir), time of division (Tdiv), length at birth (Lbir), length at division (Ldiv), cell id of the parent (PId) and cell centroid position at the time of division (Pdiv) and each cell (of one channel) has an unique cell id given by the row number in the cell data (Tab. I). The cells in the first frame of each channel is given id according to their position relative to the closed end of the channel where we refer the cell at the closed end as *old-pole cell*. Cells created from the division of these initial cells are given cell id according to their birth, a cell born earlier shall have a smaller cell id than a cell born later on. The first and seventh columns record the current index (Cind ∼ relative position of the cell in channel at current frame) and the index at the time of division (Idiv) respectively (Tab. I). These two quantities are dynamic and keep changing as the analysis progresses and these values were used in various tasks during segmentation. The current index become zero when a cell divides or exits the channel. All the lengths recorded are in the units of pixels here. The values can be decimal because the quantities are calculated from the major axis length and centroid position of the fitted ellipse. This however does not necessarily mean we can achieve sub-pixel accuracy.

**TABLE I.** Cell division data structure.

| Cind | Tbir | Tdiv | Lbir | Ldiv | PId | Idiv | Pdiv |
| --- | --- | --- | --- | --- | --- | --- | --- |
| 0.00 | 0.00 | 11.00 | 0.00 | 27.428474 | 0.00 | 1.00 | 31.048128 |
| 0.00 | 0.00 | 30.00 | 0.00 | 26.446321 | 0.00 | 3.00 | 71.430233 |
| 0.00 | 0.00 | 36.00 | 0.00 | 30.262136 | 0.00 | 5.00 | 106.938776 |
| 0.00 | 0.00 | 6.00 | 0.00 | 25.324549 | 0.00 | 4.00 | 81.472603 |
| 0.00 | 0.00 | 8.00 | 0.00 | 28.690051 | 0.00 | 6.00 | 112.565934 |
| 0.00 | 0.00 | 0.00 | 0.00 | 0.00 | 0.00 | 0.00 | 0.00 |
| 0.00 | 0.00 | 0.00 | 0.00 | 0.00 | 0.00 | 0.00 | 0.00 |
| 0.00 | 6.00 | 0.00 | 11.395244 | 0.00 | 4.00 | 0.00 | 0.00 |
| 0.00 | 6.00 | 0.00 | 13.143442 | 0.00 | 4.00 | 0.00 | 0.00 |
| 0.00 | 8.00 | 0.00 | 13.518004 | 0.00 | 5.00 | 0.00 | 0.00 |
| 0.00 | 8.00 | 0.00 | 14.304699 | 0.00 | 5.00 | 0.00 | 0.00 |
| 0.00 | 11.00 | 44.00 | 13.168872 | 28.551908 | 1.00 | 1.00 | 32.764706 |
| 0.00 | 11.00 | 44.00 | 14.444769 | 29.267566 | 1.00 | 2.00 | 59.554404 |

For the sake of more clarity here we elaborate a small segment of the cell data (Tab. I) with corresponding segmented images (Fig. 3). If we notice the time of birth column we see three divisions (at 6, 8 and 11 min) within first 15 frames (15 mins). In the first cell division the cell with cell id= 4 divides to produce two daughter cells with cell id= 8 and cell id= 9 (notice the PId for 8th and 9th row). Similarly cells with cell id= 5 and cell id= 1 divides at time *t* = 8 min and *t* = 11 min to give rise to cells with ids 10, 11, 12, 13 respectively (Fig. 3). It is important to notice that in this small segment only the cell with cell id= 13 has both non-zero birth time and division time from which a cell cycle duration can be calculated. The cells with cell id 8 to 11 exit the channels after birth (at time *t >* 15 min) before they could divide to complete one cell cycle. As the analysis progresses we gather more cells with complete cell cycle information.

**FIG. 3.**
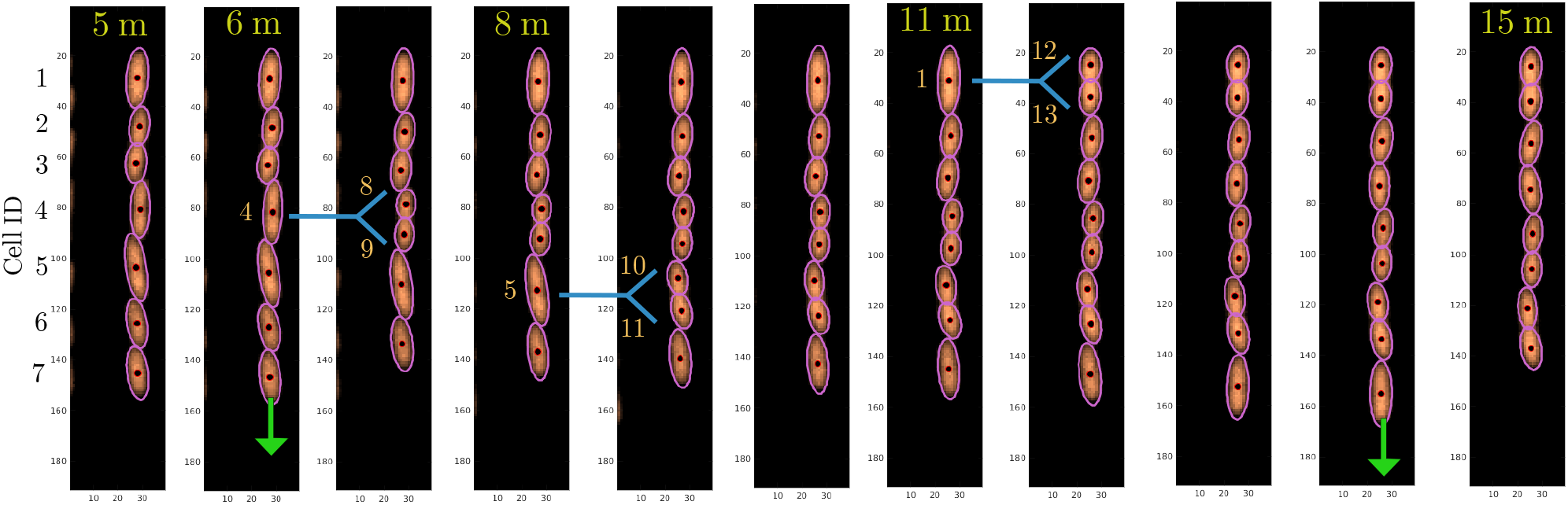
Cell division and extrusion: Example of an workflow output of 11 consecutive frames (from some time *t* = 5 min to *t* = 15 min). Cell division events (blue fork) were detected in 2nd, 4th and 7th frames and cell extrusion events (green arrow) were detected in 2nd and 10th frame.

The cell data recorded can be used to evaluate various cell division statistics. Here we present some examples of the things that can be calculated dividing them in three categories:

- The phenomenology of cell division can be probed by checking the three major classes of cell division rule, namely the adder, sizer and timer. We plot added length (Δ_*L*_), length at division (*L*_*d*_) and cell cycle duration (*τ*) with the length at birth (*L*_*b*_). The plots indicate a strong adder nature of cell division as observed in *E. coli* [14]. See Fig. 4 for details.

**FIG. 4.**
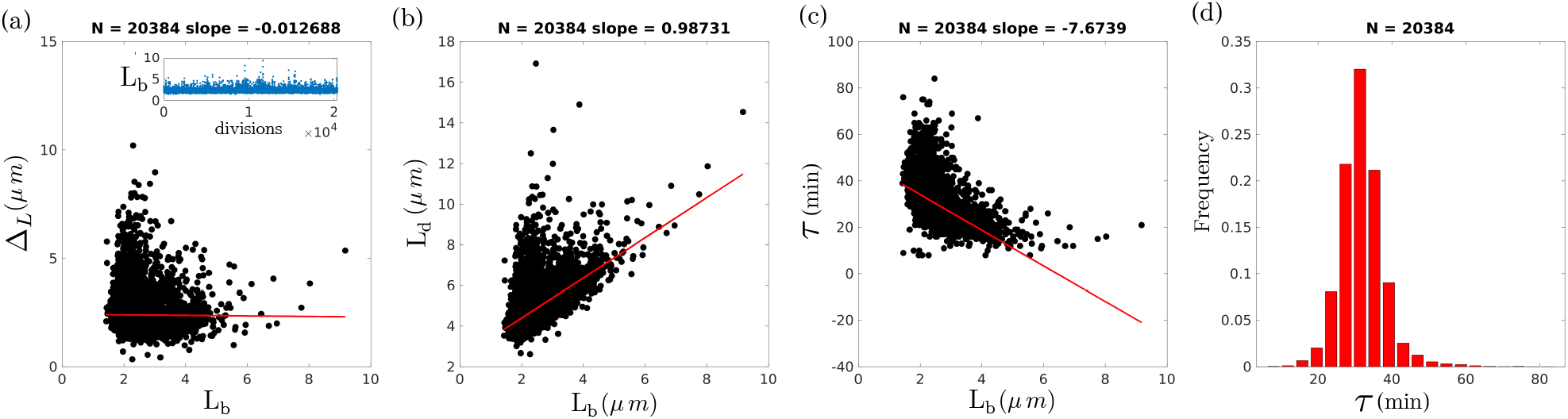
Cell division statistics: (a) We plot added length (Δ_*L*_), (b) length at division (*L*_*d*_) and (c) cell cycle duration (*τ*) with the length at birth (*L*_*b*_) and find the cell division strongly support a adder like mechanism of growth where cells add a constant volume (length, assuming with remains same) between birth and division in each cell cycle. (d) The division time distribution for all the cell divisions recorded. (inset) Birth length at each division shows the variation in the quantity. *N* denote the number of cell divisions detected.
- The features of cell division in subsequent generations can be probed from our data. Here we show the Pearson’s correlation of birth length in the first and second generation (mother and daughter divisions) and in the first and third generation (mother and granddaughter divisions). The plots show a decrease in birth length correlation with progressing generation which indicates that stochasticity (of various origin) in cell growth and division destroys correlation over the generations. See Fig. 5a-c for details.

**FIG. 5.**
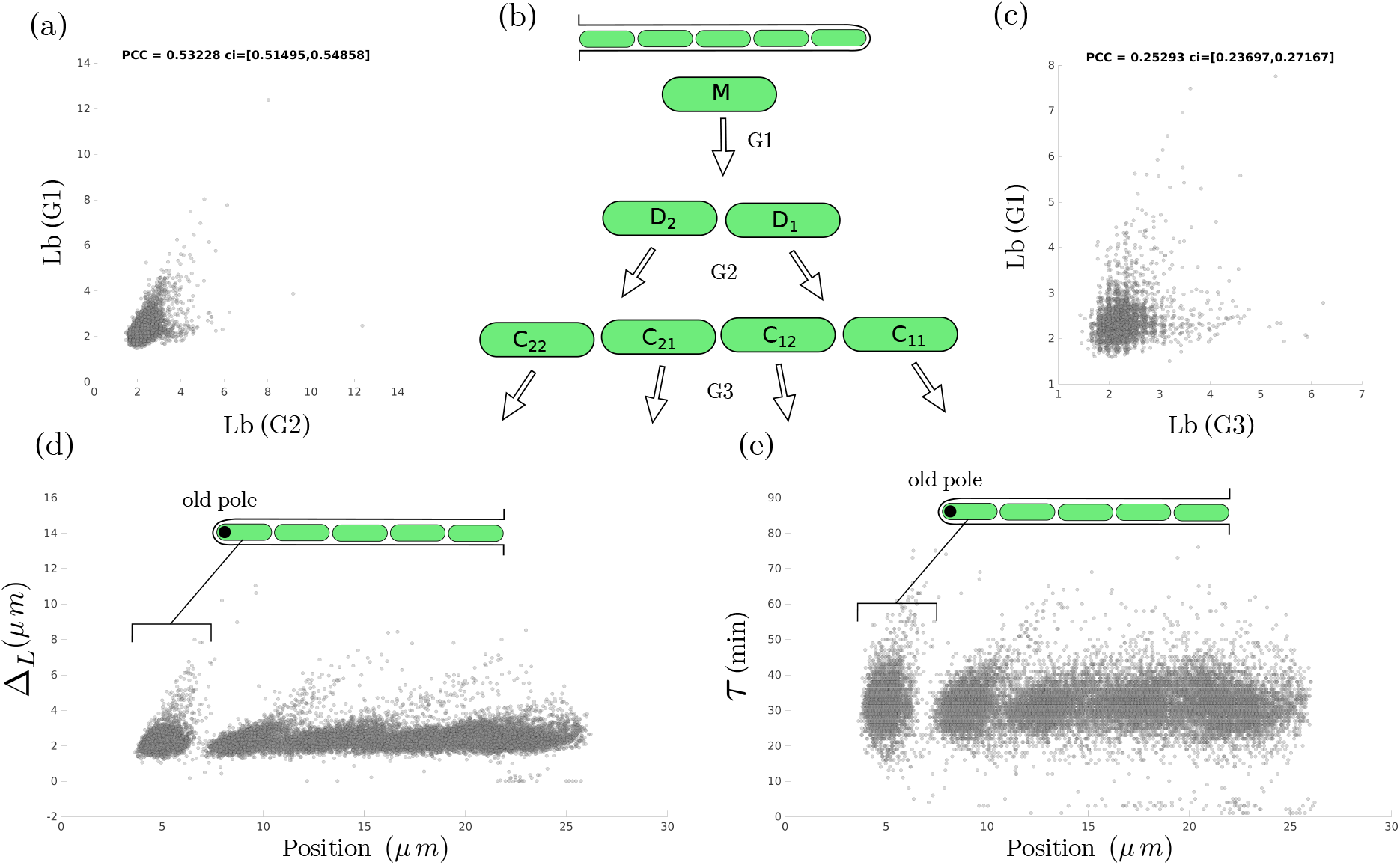
Correlations over subsequent generations and pole age: (a-c) Pearson’s correlation constant (*PCC*) of birth length in the first and second generation (mother and daughter divisions) and in the first and third generation (mother and granddaughter divisions) shows a decay in correlation with progressing generation. The confidence interval (*ci*) was calculated for a 50% window. (d-e) Cell division features were calculated with respect to the position of division (i.e., position of centroid of the dividing cell). This analysis can be useful in testing if the old cell in the end of the channel divides differently (or “ages”) than the other newly born cells. We find no significant change in division quality over the observed time window.
- Variation in cell division quantities with the position of division. We probe the added length and cell cycle duration for divisions at different position and do not find any significant variation in this particular experimental data. But the *old-pole cell* division features may deviate from the population average [15] as it does not exit the channel and hence contains the old pole (the pole closer to the channel closed end). Correlating cell division features with pole age can be useful to determine such effects if there exist any [1]. See Fig. 5d-e for details.

## V. EXPERIMENTAL METHODS

For the data presented in Fig. 1 experiments were performed on the *E. coli* strain MG1655 constitutively expressing green fluorescent protein [16]. Cells were grown in Luria Bertani medium (LB) at 37° Celsius for 3 hours before centrifugation. The pellet was resuspended in 200 *µ L* of LB which was injected into the microfluidic device, and the cells were allowed to diffuse into the growth channels. The “mother machine” was fabricated as described by Wang *et al* [1]. The device was placed in a stage-top incubator (Okolab) placed on a microscope (Olympus IX81) with a continuous supply of LB (700 *µ L/hour*), both kept at 37° Celsius. Fluorescence images (ex: 490 nm) were taken every 2 min by an EMCCD camera (Photometrics Prime).

## VI. DISCUSSION

In this document we have presented an image segmentation workflow which can be used in analysis of image data obtained from a microfluidic mother machine experiment with minimal manual supervision. Our workflow uses various local and global quantities to implement a slew of adaptive routines to minimize errors due to experimental aberrations such as cell intrusion, sticky cells at channel exit, intermittent high intensity cell clusters in flow channels etc. We have successfully analyzed data from different experiments and with different media conditions (not presented here) with very small margins of error (*<* 1% of the cell divisions captured are found to be erroneous). We hope our method will be useful to the community as an analysis tool. For more detailed documentation and current versions of the workflow please refer to the [Github repository for SAM](https://github.com/DebsankarBanerjee/Segmentation-and-analysis-of-mother-machine-data). Currently our code does not implement any machine learning methods but we shall include such methods in future versions of the workflow to lessen manual inputs in the process.

## VII. CODE AVAILABILITY

The main routines with full data analysis modules (paired with shell scripting) and test image files are available as the [Github repository](https://github.com/DebsankarBanerjee/Segmentation-and-analysis-of-mother-machine-data) of SAM. A MATLAB [console only version](https://github.com/DebsankarBanerjee/Segmentation-and-analysis-of-mother-machine-data/tree/Console-version) is also available which can easily be used in many platforms (checked for Mac, Windows and Linux). For beginners we have made a [trainer](https://github.com/DebsankarBanerjee/SAM-Trainer) with an aim to illustrate how SAM parameters change in two very different experiments. We suggest users to start with this version when they tune SAM for their data.

## ACKNOWLEDGMENTS

We thank Dr. Shashi Thutupalli for his support, useful feedback and a careful reading of the manuscript. We thank Dr. Mukund Thattai for many insightful discussions. DB thanks [Image Analyst](https://www.mathworks.com/matlabcentral/profile/authors/1343420) and numerous other contributors of MathWorks community. We acknowledge support from the Simons Foundation and NCBS-TIFR. G.S. acknowledges support from the Max Plank Society as a part of Max-Plank Partner Group to Shashi Thutupalli.

